# One hundred million years history of bornavirus infections hidden in vertebrate genomes

**DOI:** 10.1101/2020.12.02.408005

**Authors:** Junna Kawasaki, Shohei Kojima, Yahiro Mukai, Keizo Tomonaga, Masayuki Horie

## Abstract

Although viruses have threatened our ancestors for millions of years, prehistoric epidemics of viruses are largely unknown. Endogenous bornavirus-like elements (EBLs) are ancient viral sequences that have been integrated into animal genomes. These elements can be used as molecular fossil records to trace past bornaviral infections. In this study, we systematically identified EBLs in vertebrate genomes and revealed the history of bornavirus infections over nearly 100 million years. We found that ancient bornaviral infections have occurred in diverse vertebrate lineages, especially in primate ancestors. Phylogenetic analyses indicated that primate ancestors had been infected with various bornaviral lineages during evolution. Moreover, EBLs in primate genomes formed clades according to their integration ages, suggesting that epidemic lineages of bornaviruses had changed chronologically. However, we found that some bornaviral lineages coexisted with primate ancestors and underwent repeated endogenizations for tens of millions of years. Furthermore, this viral lineage that coexisted with primate ancestors was also endogenized in some ancestral bats. Notably, the geographic distributions of these bat ancestors have been reported to overlap with the migration route of primate ancestors, suggesting that long-term virus-host coexistence could have expanded the geographic distributions of the viral lineage and might have spread their infections to new hosts. Thus, our findings describe hidden virus-host co-evolutionary history over geological timescales, including chronological change in epidemic bornaviral lineages, long-term virus-host coexistence, and expansion of viral infections.

## Introduction

Viral infectious diseases profoundly affect human health, livestock productivity, and ecosystem diversity. Similar to recent viral outbreaks [1], our ancestors were also probably challenged by viral epidemics. Investigations of past viruses using historical specimens have provided insights into the origin and spread of viral infections [2–6]. However, the epidemic history of viruses across hundreds of millions of years is largely unclear.

Endogenous viral elements (EVEs) are formed by the occasional integration of ancient viral sequences into the host germline genomes [7]. EVEs are millions of years old and provide critical information on ancient viruses, such as their host ranges [8, 9], evolutionary timescales [10], or geographical distributions [11]. Endogenous bornavirus-like elements (EBLs) are the most abundant viral fossils of RNA viruses found in vertebrate genomes [12, 13]. Therefore, EBLs could help us trace the epidemic history of bornaviral infections on geological timescales and are good model systems to study long-term virus-host co-evolutionary history.

The family of *Bornaviridae* consists of three genera, *Orthobornavirus, Carbovirus,* and *Cultervirus* [14]. Until 2018, only the genus *Bornavirus* (today *Orthobornavirus*), which includes viruses that cause immune-mediated neurological diseases in mammals and birds, constituted the family *Bornaviridae* [15]. However, two new genera, *Carbovirus* and *Cultervirus*, were established upon discovering novel bornaviral species in carpet pythons and sharpbelly fish samples, respectively [16, 17]. Furthermore, these discoveries also led to the identification of novel EBLs classified into these bornaviral genera [16]. Since most previous studies have exclusively used orthobornaviruses for EBL detection [10, 12, 13, 18–20], numerous EBLs may remain to be detected and analyzed. Therefore, current understanding of the epidemic history of ancient bornaviruses is probably incomplete.

In this study, we searched for EBLs derived from three bornaviral genera and characterized the evolutionary timescale, host range, geographical distribution, and genetic diversity of ancient bornaviruses to reconstruct the long-term history of their infections. Large-scale dating analysis revealed that ancient bornaviral infections have occurred in diverse vertebrate lineages for nearly 100 million years. Primate ancestors, in particular, had been repeatedly infected with ancient bornaviruses. Phylogenetic analyses of EBLs in primate genomes showed clustering according to their integration ages, suggesting that epidemic lineages of bornaviruses in primate ancestors have changed chronologically. Furthermore, some bornaviral lineages may have coexisted with primate ancestors for tens of millions of years. Interestingly, we found that long-term virus-host coexistence could have expanded the geographic distributions of the virus and generated new infections in some bat ancestors. Thus, our findings describe virus-host coevolutionary history over geological timescales, which cannot be deduced from research using extant viruses.

## Results

### Systematic identification of EBLs in the host genomes

To systematically identify EBLs, we searched for bornavirus-like sequences in the genomic data of 969 eukaryotic species by tBLASTn using bornaviral protein sequences from all genera in the family *Bornaviridae* as queries (**Fig 1A**). Next, we concatenated these sequences based on their location in the host genome and their alignment positions to extant bornaviral proteins because most bornavirus-like sequences were fragmented due to mutation after endogenization.

**Fig. 1.**
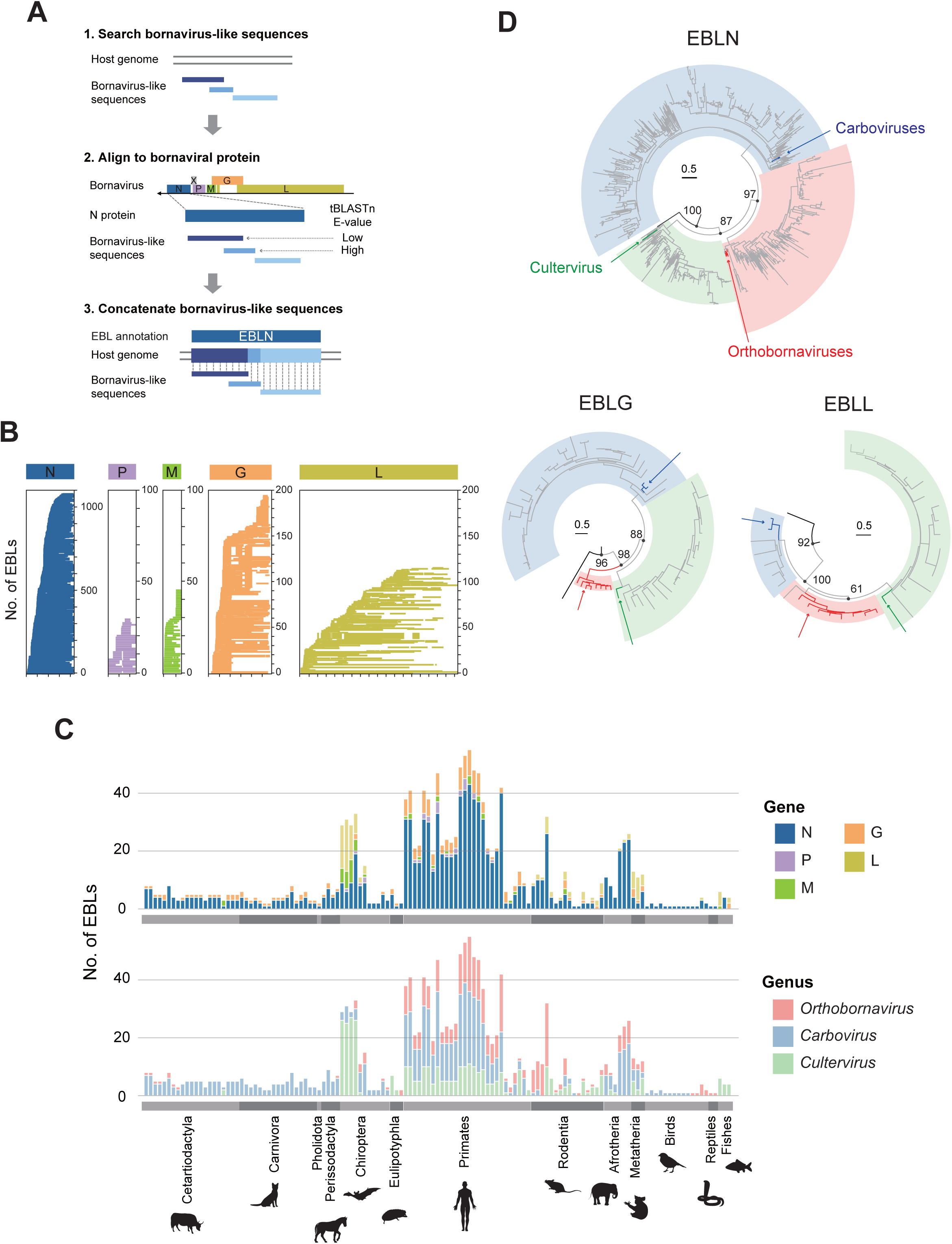
Identification of EBLs in the vertebrate genomes. **(A)** Schematic diagram for the procedure to identify EBLs. First, bornavirus-like sequences were detected from host genomes by tBLASTn using extant bornaviral sequences as queries. Second, the detected bornavirus-like sequences were aligned with corresponding proteins of extant bornaviruses. Third, the bornavirus-like sequences were concatenated based on the host genomic locations and alignment positions with the bornaviral proteins. When several bornavirus-like sequences were detected in the same genomic positions, the sequence with higher reliability (low E-value score in the tBLASTn search) was used for EBL sequence reconstruction. **(B)** Alignment coverage plot of EBLs. The scales on the x-axis in the alignment are marked at intervals of 100 amino acids. The y-axis indicates the number of EBLs identified in this study. **(C)** Numbers of EBLs in the host genomes. The x-axis indicates the vertebrate species and the y-axis indicates the number of EBLs identified in the species genome. The bar color shows the bornaviral gene (upper panel) or the genus from which the EBL originated (lower panel). **(D)** Phylogenetic trees of EBLs and extant bornaviruses. These trees were constructed by the maximum likelihood method using the amino acid sequences of EBLs and extant bornaviral proteins. The branch colors indicate the sequence groups: EBLs (gray), extant nyamivirus used as outgroup (black), extant orthobornaviruses (red), extant carboviruses (blue), and extant cultervirus (green). Colored arrows mark extant bornaviruses. The highlights correspond to the current bornaviral classifications: genus Orthobornavirus (light red), genus Carbovirus (light blue), and genus Cultervirus(light green). Representative supporting values (%) are shown on branches. The scale bars indicate genetic distances (substitutions per site).

The bornaviral genome encodes 6 viral proteins: nucleoprotein (N), phosphoprotein (P), matrix protein (M), envelope glycoprotein (G), large RNA-dependent RNA polymerase (L), and accessory protein (X) [21]. EBLs are considered to have been formed by integrations of bornaviral mRNA into the host genome; EBLs derived from N, M, G, and L genes, designated EBLN, EBLM, EBLG, and EBLL, respectively, have been reported so far [13]. Our EBL search identified 1,465 EBLs in 131 vertebrate species, including 1,079 EBLNs, 30 EBLPs, 46 EBLMs, 195 EBLGs, and 115 EBLLs (**Fig 1B and 1C**). Notably, we identified EBLPs in animal genomes for the first time, although these were considered difficult to detect due to methodological limitations, such as low sequence conservation of P genes among bornaviruses [22].

We next classified ancient bornaviruses from which EBLs originated. Since some EBLs were too short to construct reliable phylogenetic trees, we sought to classify all the EBLs based on sequence similarity scores in the tBLASTn search. However, if there is an unknown genus consisting exclusively of ancient bornaviruses, the method based on sequence similarity with modern viruses may lead to misclassification. To assess this possibility, we first performed phylogenetic analyses using relatively long EBLs and their gene counterparts in modern bornaviruses. **Fig 1D** shows that EBLs were clearly divided into three clades corresponding to the current bornaviral classification [14]. Therefore, we decided to apply the sequence similarity-based method for classifying all the EBLs into current bornaviral genera (**details in Materials and Methods**). Based on the similarity scores, the EBLs were classified into 364 orthobornaviral, 729 carboviral, and 372 culterviral EBLs (**Fig 1C**). Among 1465 EBLs, 870 loci were undetectable by a search using orthobornaviral sequences that have been almost exclusively used for EBL detection as queries. Therefore, the EBL search found numerous previously undetected loci and created a comprehensive dataset for reconstructing the history of bornavirus infections.

### Large-scale dating analysis for bornaviral integration ages

EVE integration ages can be estimated based on the gene orthology [7], and thus we sought to determine the presence and absence patterns of orthologous EBLs. Here, we developed a network-based method to handle the large datasets (**S1 Fig**). Briefly, we first constructed an all-against-all matrix of alignment coverages by pairwise sequence comparison among EBL integration sites. Next, we constructed a sequence similarity network using the matrix and extracted community structures from the network in order to divide the EBLs into groups based on their orthologous relationships. Finally, we manually checked the groupings to avoid inaccurate estimates (**details in Materials and Method and S2 Fig**).

We divided 1,465 EBLs into 281 groups by our network-based dating method (**S1 Table**). These groupings reflected the alignment coverage among EBL integration sites (**S1 Fig**). We divided these groups into two categories: 113 groups of “EBLs with orthologs” and 168 groups of “EBLs without orthologs” (**Fig 2A**). “EBLs with orthologs” share the same bornaviral integration in each group, while each group of “EBLs without orthologs” consists of a single EBL locus. Such “EBLs without orthologs” might be young integrations that occurred after the divergence of hosts from their sister species. Alternatively, the lack of orthologs may simply be a methodological limitation due to the inaccessibility of the genomic data of sister species. For example, only three distantly related species of Eulipotyphla, in which no EBL orthologous relationships could be detected, were present in our database (**Fig 2A**). This suggests that accumulating genomic data could help estimate integration ages with higher accuracy.

**Fig. 2.**
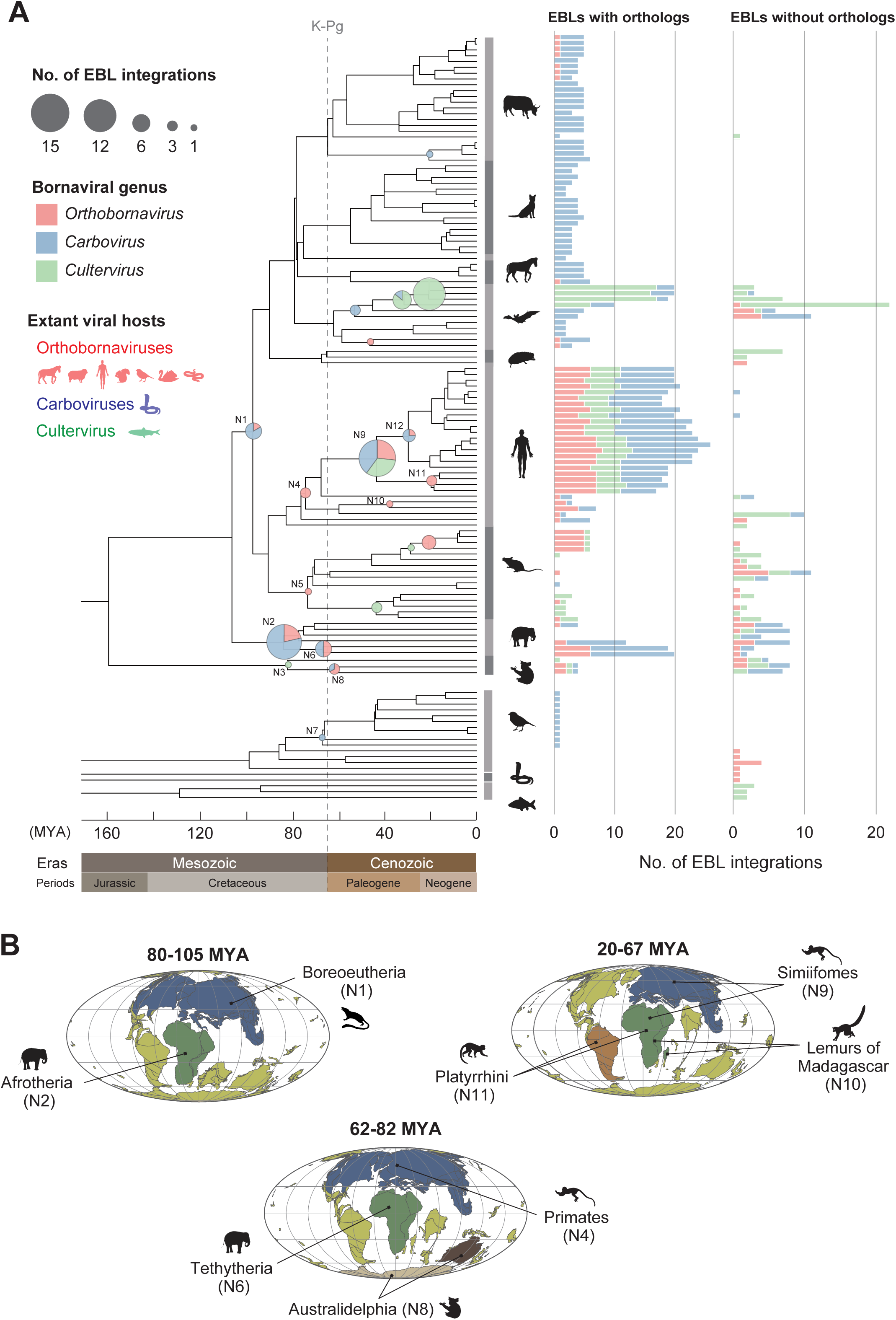
The history of bornaviral integration events for approximately 100 million years. **(A)** Bornaviral integration events during vertebrate evolution. The evolutionary tree of vertebrates was obtained from the TimeTree database. The positions of pie-charts on the tree indicate the lower limit ages of bornaviral integration events, and their size shows the number of events in each period determined based on the gene orthology. Annotations in the internal nodes on the tree indicate the common ancestors of Boreoeutheria (N1), Afrotheria (N2), Metatheria (N3), Primates (N4), Rodentia (N5), Tethytheria (N6), Passeriformes (N7), Australidelphia (N8), Simiiformes (N9), Lemuroidea (N10), Platyrrhini (N11), and Catarrhini (N12). The representative hosts of extant bornaviruses are shown by animal silhouettes on the left side of the tree. The right panel shows the numbers of EBLs categorized into EBLs with orthologs or EBLs without orthologs in each species. The definitions of these categories are described in the section titled Large-scale dating analysis for bornaviral integration ages. The bar colors show the viral genus as indicated on the left side of the tree. **(B)** A schematic diagram of the geographical distributions of ancient bornaviruses and their hosts in each era. The colored continents, except for yellow, indicate the continents where bornaviral endogenization may have occurred: Laurasia or Eurasia (blue), Africa (green), Antarctica (beige), Australia (dark brown), and South America (brown). The biogeography of hosts during their evolution was cited from previous reports (**S2 Table**). The plate tectonic maps were downloaded from ODSN Plate Tectonic Reconstruction Service (http://www.odsn.de/odsn/services/paleomap/paleomap.html).

### Bornaviral infections have occurred since the Mesozoic era

Our dating analysis allowed us to trace the history of bornavirus infections back to 100 million years ago (MYA) (**Fig 2A**). The oldest records of bornaviral infection have been reported in ancestral afrotherians at least 83.3 MYA [10, 20]. Here, we found six EBLs that were orthologous among species of Boreoeutheria, suggesting that the oldest bornavirus infections had occurred at least 96.5 MYA. Additionally, we identified 18 bornaviral integration events in the Mesozoic era, which occurred in the ancestors of Afrotheria, Tethytheria, Metatheria, Primates, and Rodentia. Besides, we found the first record of bornavirus infection in Mesozoic birds in the ancestor of Passeriformes at least 66.6 MYA. Thus, these results provide strong evidence that bornaviral infections had already occurred in multiple vertebrate lineages in the Mesozoic era.

### Ancient bornaviral infections in various vertebrate lineages

We found that ancient bornaviruses infected much broader vertebrate lineages than modern bornaviruses are known to infect (**Fig 2A**). Modern orthobornavirus infections have been reported in ungulate animals, shrews, squirrels, humans, a wide range of birds, and garter snakes [15]. However, we identified multiple endogenizations of ancient orthobornaviruses in mice, afrotherians, and marsupials, which have not been reported as host species of modern orthobornaviruses (**Fig 2A**). In particular, a previous survey of bornaviral reservoirs did not detect orthobornavirus infections in mice [23]. Furthermore, the extant carboviruses and cultervirus were detected only in carpet pythons and sharpbelly fish, respectively [16, 17]. In contrast, ancient viruses belonging to these genera endogenized in various host lineages, including mammals and birds (**Fig 2A**). These results indicate that ancient bornaviruses infected a wider range of vertebrate lineages than known extant bornaviruses.

### The geographical distributions of ancient bornaviral infections

We performed integrative analysis of bornaviral endogenizations and mammalian biogeography dynamics to infer the geographical distributions of ancient bornavirus infections (**S2 Table**). Our results suggest that ancient bornavirus infections occurred in different continents: Laurasia and Africa in the Mesozoic era, Antarctica or Australia around the K-Pg boundary, and possibly Eurasia, Africa, or South America in the Cenozoic era (**Fig 2B**).

First, we identified EBLs in animals that inhabited Laurasia and Africa in the Mesozoic era (**N1, N2, N4, and N6 in Fig 2B**). It has been reported that ancestors of Boreoeutheria and Primates were distributed in Laurasia [24–27], while those of Afrotheria and Tethytheria were found in Africa [24, 25, 28]. These results suggest that bornaviral infections might have spread in Laurasia and Africa during the Cretaceous period. Second, we identified EBLs integrated into the genome of Australidelphia ancestors, but not in other marsupials in South America (**N8 in Fig 2**). Since ancestral Australian marsupials are considered to have moved from South America to Australia via Antarctica [29, 30], bornavirus infections are likely to have occurred in Antarctica or Australia.

Furthermore, we identified bornaviral endogenizations that occurred in ancestral primates in the Cenozoic era. First, we found EBLs integrated into the genome of the ancestor of the Madagascar lemur (**N10 in Fig 2**); however, EBLs were not identified in African galagos, suggesting that bornavirus infections occurred in Madagascar Island. On the other hand, EBL integration age was estimated at 37.8-59.3 MYA, which overlapped with the migration of lemur ancestors from Africa to Madagascar Island around 50-60 MYA [31]. This overlap presents an alternate possibility that bornaviral endogenization had occurred in African animals before they migrated to Madagascar. Second, we identified several EBLs integrated into the ancestral Platyrrhini genomes (**N11 in Fig 2**). Ancestors of Platyrrhini are presumed to have migrated from Africa to South America during their divergence from Simiiformes [26, 32], thus providing evidence of bornavirus infections in these continents (**see Discussion**). Taken together, these results suggest worldwide occurrence of ancient bornavirus infections.

### The complex history of bornaviral infections during primate evolution: epidemics of distinct bornaviral lineages in each age

We found that bornaviruses in the three genera repeatedly endogenized during primate evolution (**Fig 2A**). We inferred phylogenetic relationships among ancient bornaviruses in each genus using EBLNs that are the most abundant records among EBLs (**Fig 3A-C and S3 Fig**) in order to understand the origin of endogenizations of bornaviral lineages into primate ancestor genomes.

**Fig. 3.**
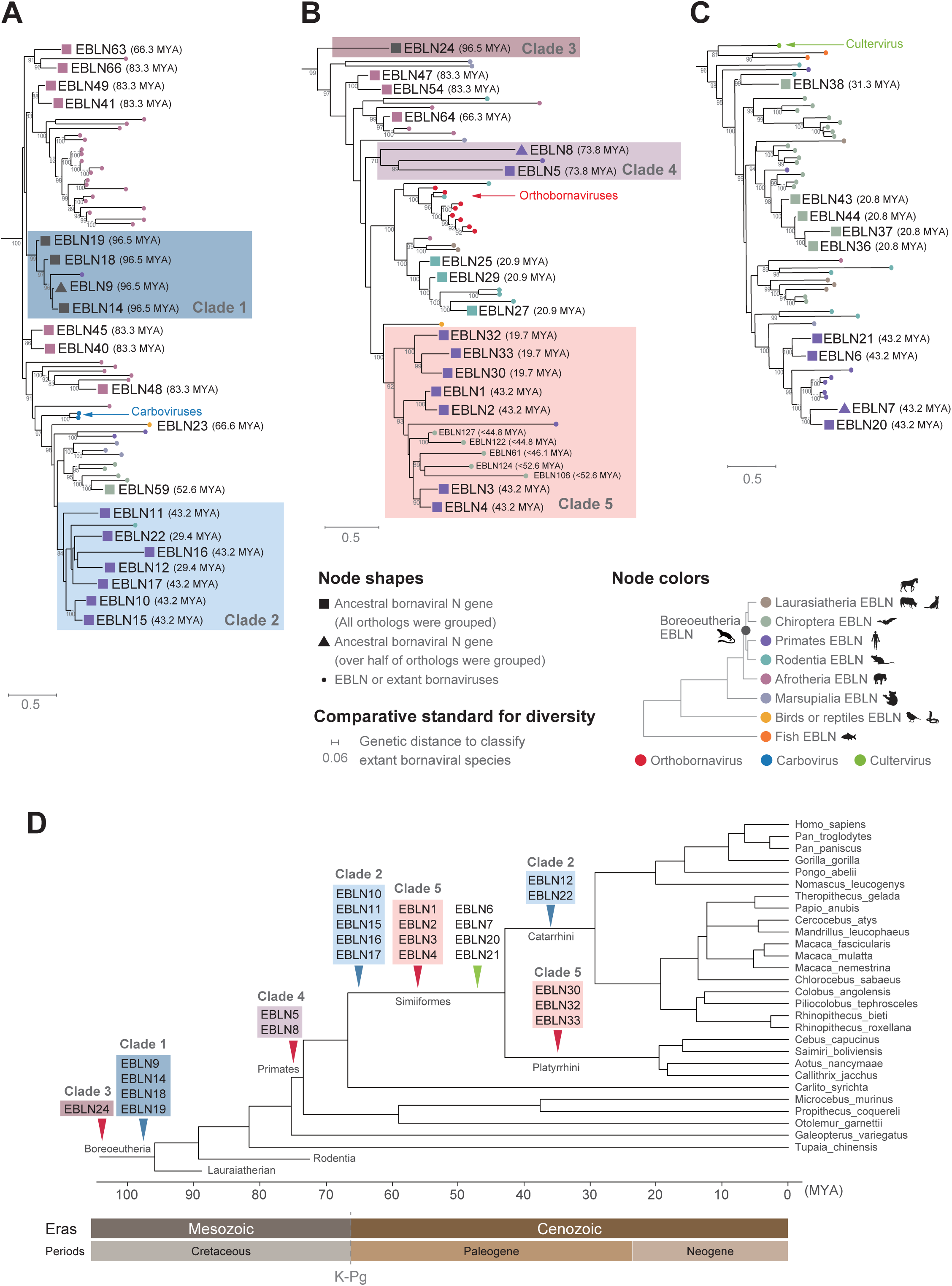
Phylogenetic relationships of ancient bornaviruses that infected primate ancestors. **(A-C)** Phylogenetic analyses of ancient and modern bornaviral N genes. These trees were constructed by the maximum likelihood method using the amino acid sequences of EBLNs and extant bornaviral N proteins of genus Carbovirus (A), Orthobornavirus (B), or Cultervirus (C). Colored arrows mark extant bornaviruses. The square and triangle nodes indicate collapsed clades containing all and over half of the orthologs used in the phylogenetic analyses, respectively. Phylogenetic trees with all expanding nodes are available in S3 Fig. The node colors indicate the host lineages of ancient bornaviruses or extant bornaviral genera as indicated in the lower right panel. The colored boxes highlight the bornaviral lineages endogenized during primate evolution. The number on the branches are bootstrap values (%) based on 1000 replications. The scale bars show genetic distances (substitutions per site). The genetic distance to distinguish extant bornaviral species is shown as the comparative standard for estimating the genetic diversity of ancient bornaviruses. **(D)** EBLN integration events during primate evolution. Arrowheads indicate the occurrence of ancient bornaviral integrations: orthobornaviral EBLN (red), carboviral EBLN (blue), and culterviral EBLN (green). The colors of highlighted boxes correspond to ancient bornaviral lineages shown in (A-C).

Consequently, we found several distinct lineages of bornaviruses that had sequentially endogenized during primate evolution, rather than a single bornaviral lineage that had repeatedly endogenized. For example, the carboviral EBLNs in the primate genome were clearly divided into two viral lineages: the clade 1 viral lineages endogenized in Boreoeutherian ancestors and the clade 2 viral lineages endogenized in Simiiformes and Catarrhini ancestors (**Fig 3A and 3D**). Furthermore, orthobornaviral EBLNs formed three different clades according to their integration ages (**clades 3 to 5 in Fig 3B and 3D**). These results suggest that different bornaviral lineages were prevalent during primate evolution across different eras.

Next, to infer how diverse bornaviruses have endogenized during primate evolution, we calculated the genetic distances between these ancient viral lineages (clades 1 to 5) in our phylogenetic tree (**S3 Table**). Using genetic distance as a comparative standard for classifying extant species of bornaviruses, we found that the genetic diversity among these ancient bornaviral lineages was higher than that among extant bornaviral species (**Fig 3A-C and S3 Table**). Thus, we infer that recurrent bornaviral endogenizations during primate evolution occurred due to infections of multiple bornaviral lineages comparable to different viral species.

### The complex history of bornaviral infections during primate evolution: long-term virus-host coexistence

In addition to the sequential infections of primate ancestors by distinct bornavirus lineages (**Fig 3**), we found that some lineages might have established long-term coexistence with the hosts. For example, the clade 2 carboviral lineage has repeatedly endogenized in Simiiformes and Catarrhini ancestors between 29.4 and 67.1 MYA (**Fig 3A and 3D**). Furthermore, endogenizations of the clade 5 orthobornaviral lineage have recurred in Simiiformes and Platyrrhini ancestors between 19.7 and 67.1 MYA (**Fig 3B and 3D**). These results suggested that these bornaviral lineages have coexisted with primate ancestors for tens of millions of years. In summary, we described the complex history of recurrent bornaviral endogenizations during primate evolution, including sequential infections of diverse bornaviral lineages and long-term virus-host coexistence.

## Discussion

Snapshots of ancient bornaviral infections have been reported since the discovery of EBLs [10, 12, 13, 16, 18–20]; however, the long-term history of bornavirus infections has remained unclear. Here, we systematically identified EBLs in 131 vertebrate species (**Fig 1**) and reconstructed the epidemic history of bornaviral infections for approximately 100 million years (**Fig 2**). To our knowledge, this is the first report to comprehensively trace the history of RNA virus infections over geological timescales. Furthermore, phylogenetic analyses suggested differences in the epidemic lineages of bornaviruses during primate evolution across different geological ages as well as coexistence of some lineages with ancestral primate ancestors for tens of millions of years (**Fig 3**). Virus-host co-divergence alone, which is thought to be as the background of viral evolutionary history [17], is insufficient to explain this mixed pattern. Therefore, our findings suggested that the virus-host coevolutionary relationships had been dramatically changed over geological timescales, which complicated the viral evolutionary history.

We also found that various host lineages might have been infected by phylogenetically related bornaviruses in each geological age (**Fig 3**). Although EBLs are the most abundant RNA virus fossils, it is not easy to trace the details of viral transmission because EBLs have only rarely been fossilized. Nonetheless, our phylogenetic analyses showed that bornaviruses closely related to the lineage that caused epidemics in ancestral primates had almost contemporaneously endogenized in the other animals (**Fig 3 and S3 Fig**). For example, carboviruses similar to the clade 1 viral lineage had also endogenized in ancestral afrotherians around the late Mesozoic era (**EBLN41, 49, 63, and 66 in Fig 3A**). Additionally, carboviruses similar to the clade 2 lineage endogenized in ancestors of Yangochiroptera bats from the late Mesozoic to Cenozoic era (**EBLN59 in Fig 3A**). A similar tendency was observed in the orthobornaviral phylogenetic tree and extant viruses as well: genetically similar orthobornaviruses infect various host species in mammals, birds, and reptiles at present (**Fig 3B**). These results suggest that bornaviral lineages have spread to various hosts in each era, and epidemic lineages of bornaviruses have changed over time.

By integrating information on host geographical distributions and phylogeny of bornaviruses, we found long-term coexistence of ancient bornaviruses with primate ancestors that could have expanded the viral lineage to other continents. **Fig 3B** shows that the clade 5 orthobornaviral lineage has repeatedly endogenized in ancestors of Simiiformes and Platyrrhini. Ancestors of Simiiformes were reportedly distributed in Eurasia or Africa, while Platyrrhini ancestors likely migrated from Africa to South America during their divergence from Simiiformes (**Fig 2B**) [26, 32]. These results suggested that the clade 5 viral lineage could have moved between the continents along with host migrations. Interestingly, viruses in the clade 5 lineage also endogenized in several bat genomes (**Fig 3B**), such as Rhinolophus bats (EBLN61), Desmodus bats (EBLN106 and EBLN124), and Miniopterus bats (EBLN122 and EBLN127). Since Rhinolophus and Desmodus bats have reportedly originated in Eurasia and South America, respectively [33], the clade 5 viral lineage may have moved across continents along with primate migrations and been further transmitted to these bat ancestors. Thus, long-term virus-host coexistence could have expanded the viral geographic distributions and generated new infections in other hosts.

We found that the number of bornaviral integration events varied according to the host lineage (**Fig 2A**). In non-mammalian vertebrates, we observed low frequencies of bornaviral integrations, consistent with previous studies [34, 35]. Furthermore, the numbers of bornaviral endogenizations differ among descendant lineages that diverged from boreoeutherian ancestors. Remarkably, bornaviral endogenizations rarely occurred in most of the laurasiatherian lineages, but repeatedly occurred in some other host lineages, including Primates, Chiroptera, and Rodentia. Factors such as viral infection, germ-line integration, and inheritance may be responsible for these differences. Further studies on the impact of these factors on EBL fixations are necessary to explain such differences.

This study also proposed that a large number of EVEs derived from unknown ancient viruses may be hidden in the host genomic data. One of the problems with paleovirological research is the methodological limitation for identifying viral fossil records comprehensively. Because the method to identify EVEs involves searching for virus-like sequences in the host genome using modern viral sequences as queries, the detectability of EVEs depends on sequence similarities with extant viruses. Here, we demonstrated that the EBL search using genetically diverse extant bornaviruses provided a large dataset, including previously undetectable loci (**Fig 1**). Therefore, further elucidation of extant viral diversity could help in clarifying ancient viral diversity.

Furthermore, the sequence data of EBLs could be a useful resource for exploring extant viral diversity because metagenomic analyses to detect viral infections also rely on sequence similarities with known viral sequences. Our phylogenetic analyses indicated that EBLNs originated from diverse ancient bornaviruses, and almost all EBLNs formed clades completely different from modern ones (**Fig 3 and S3 Fig**). Hence, ancient bornaviruses appear highly divergent and phylogenetically distinct from known modern viruses. Furthermore, these results raise a fascinating question regarding whether viruses genetically similar to EBLs are extinct or just yet to be discovered. Future viral metagenomic analyses using EBL sequence data may address this question. Therefore, reusing data between metagenomic analyses for extant viruses and paleovirological investigations could elucidate viral diversity and connect modern with ancient viral evolution.

In conclusion, we traced bornaviral infections over geological timescales and depicted the epidemic history of bornaviruses during vertebrate evolution. Our findings provide novel insights into coevolutionary history between viruses and hosts, which cannot be deduced from research using extant viruses.

## Materials and Methods

### Identification of EBLs in vertebrate genomic data

EBLs were identified by: (1) searching for bornavirus-like sequences in genomes of 969 eukaryotic species, (2) reconstructing EBL sequences, and (3) validating whether these EBL sequences were derived from ancient bornaviruses or not.

First, bornavirus-like sequences were screened in Refseq genomic database (version: 20190329) provided from NCBI [36] by tBLASTn (version 2.6.0+) [37] with the option “-evalue 0.1” using sequences in all genus of Bornaviridae as queries (**S4 Table**). Second, because most EBLs were detected as fragmented sequences due to mutations occurring after integration, we reconstructed EBL sequences by concatenating sequences if the following conditions were met: (1) detected bornavirus-like sequences were located within 1000 bp (EBLN, EBLP, EBLM, and EBLG) or 2000 bp (EBLL), and (2) the order of sequences in the alignment with extant bornaviral proteins were consistent with those in the host genome (**Fig 1A**). When more than two bornavirus-like sequences were detected in the same genomic position, we preferentially used the sequence with higher reliability (low E-value in the tBLASTn search). The alignment of bornavirus-like sequences and modern bornaviral proteins was conducted using MAFFT (version 7.427) with options “--addfragment” and “--keeplengths” [38]. Finally, we checked the origin of the EBL candidates based on the bit score obtained from BLASTP (version 2.9.0+) using the Refseq protein database (version: 20200313) and a database consisting of bornaviral protein sequences listed in **S4 Table**. If the candidate was more similar to the host proteins than published EBLs or other viral proteins, we considered the sequence a false positive and removed it from the analysis. After this process, only one EBLL candidate was identified in the insect genome, but we excluded this sequence in subsequent analyses. The concatenation of bornavirus-like sequences yielded over 800 EBL loci equivalent to more than half the length of the intact bornaviral proteins (**Fig 1B**).

### Dating analysis for the integrated age of EBLs

To determine orthologous relationships among EBLs, we sought to cluster loci based on the alignment coverages in pairwise sequence comparison between EBL integration sites (**S1 Fig**). First, we extracted the upstream and downstream sequences of EBLs with lengths of 15 kbp for EBLLs and 10 kbp for other EBLs. These sequences were trimmed by removing repetitive elements using RepeatMasker (version open-4.0.9) (http://repeatmasker.org) with option “-q xsmall -a -species” and RepBase RepeatMasker libraries (version 20181026) [39]. Second, we performed the pairwise alignment of these sequences using BLASTN (version 2.9.0+) and constructed an all-against-all matrix for alignment coverage among EBL integration sites. The sequence similarity network was constructed by connecting nodes when their sequence alignment coverage was over 9.0% of the flanking sequence length. Selection of the best criteria to construct a sequence network is described in the next section. The groups were extracted by detecting a community structure using Louvain heuristics. These network analyses were performed using NetworkX [40].

We simultaneously checked the phylogenetic relationships of host species with sequence alignment coverage to correctly estimate EBL ages. The contamination of sequences unrelated to true orthologous relationships leads to overestimation of integration ages, as shown in **example 4 in S4 Fig**. To avoid such issues, when multiple EBL loci were present in the same species genome and alignment coverage was lower than 50%, we considered these loci as located in different genomic sites and divided them into different groups. Furthermore, the integration ages of older elements tend to be underestimated because the alignment quality among their integration sites may deteriorate due to the accumulation of sequence changes, such as genomic rearrangement (**example 3 in S4 Fig**). Thus, by checking the phylogenetic relationships of host species and sequence alignment coverage, we combined some groups into EBLG2, EBLL2, EBLL35, or EBLL36 (**S1 Fig and S2 Fig**). For example, the EBLG2 group was previously reported to have endogenized into the genome of laurasiatherian ancestors at least 77.0 MYA [16]. This group was divided into two groups in the initial analysis, including primate and laurasiatherian loci. However, these groups were connected by low alignment coverage, which led to another hypothesis that these sequences are descendants of the same integration event in the boreoeutherian ancestor (**S1 Fig**). To test this hypothesis, we confirmed the alignment quality among the EBL integration sites using AliTV [41–43]. We found that over 70% sequence similarity covered more than 40% of the alignment by lastz (version 1.04.00) [42] with options “--noytrim, --gapped, and --strand=both” (**S2 Fig**). Therefore, we combined these groups into the same group. The cases of EBLL2 and EBLL35 were similar to that of EBLG2 (**S2 Fig**). EBLL36 contained tandemly repeated loci at close genomic locations (**S1 Table**) because we could not distinguish whether these loci were derived from independent integration events or gene duplications post-integration. Presently, we considered these loci as descendants from the same integration event to avoid overestimating the number of EBL integration events.

After curation, the dates of EBL integration events were assigned according to a vertebrate evolutionary tree provided from the TimeTree database [44]. Each EBL locus was named according to the nomenclature for endogenous retroviruses [45] (**S1 Table**). It should be noted that the number of bornaviral integration events was less than that of EBL loci shown in **Fig 1C** because redundant sequences in the genomic database used for the EBL search were grouped as the same integration event.

### Validation of the network-based dating method using human transposable elements

To validate our dating method, we compared the integration ages of human transposable elements (TEs) estimated based on genomic alignment and those estimated based on network analysis (**S4 Fig**). The genomic positions of all human TEs were obtained from the RepeatMasker database (http://www.repeatmasker.org). First, the orthologs of all human TEs were determined in 18 mammalian genomes in LiftOver (version 357) with the option “-minMatch=0.5” using genomic alignments provided by the University of California Santa Cruz (UCSC) genome browser [46]. Their integration ages were determined by the presence and absence patterns of orthologs. Second, to examine the ortholog detection rate by the network-based dating method for each timescale, we prepared test datasets by random sampling of 100 loci for each timescale based on the dating results using genomic alignment (**S4 Fig**). The details of the network-based dating method are described in the previous section. The estimation of human TE integration ages in the test datasets followed the same strategy, except that: (1) the flanked sequence of human TEs were extracted with lengths of 10 kbp from the soft-masked assembly sequence provided by the UCSC genome browser, and (2) the UCSC genome browser procedure detected repetitive elements. Finally, we compared the results between the two methods by checking the following points: predicted ages and detected orthologs (**examples are shown in S4 Fig**).

Furthermore, we tried nine different criteria to connect nodes in the sequence similarity network (**S4 Fig**) and decided to connect network edges if their sequence alignment coverage was over 9.0% of the flanking sequence length (**S4 Fig**). The concordant rates between the two methods in predicting ages of Cenozoic TEs were 88.0-100.0% according to this criterion, and those of older TEs integrated in the Cretaceous period were 43.0-66.0% (**S4 Fig**). The chain files used for LiftOver and the genome assembly sequences are listed in **S5 Table**.

### Phylogenetic analysis

We used amino acid sequences of EBLs with lengths longer than 200 amino acids for EBLL and 100 for EBLN and EBLG. Multiple sequence alignments (MSAs) were constructed by MAFFT with options “--addfragment” and “--keeplengths.” MSAs for EBL classification (**Fig 1D**) were trimmed by excluding sites where over 30% of sequences were gaps, subsequently removing sequences with less than 70% of the total alignment sites. MSA for the EBLN tree (**Fig 3 and S3 Fig**) was trimmed by excluding sites where over 20% of sequences were gaps, and subsequently removing sequences with less than 80% of the total alignment sites. Phylogenetic trees were constructed by the maximum likelihood method using IQTREE (version 1.6.12) [47]. The substitution models were selected based on the Bayesian information criterion score provided by ModelFinder [48]: VT+F+G4 for EBLNs, VT+F+G4 for EBLGs, VT+F+R3 for EBLLs (**Fig 1D**), and JTT+F+G4 for EBLNs (**Fig 3 and S3 Fig**). The branch supports were measured as the ultrafast bootstrap values given by UFBoot2 [49] with 1000 replicates. The extant viral sequences used for the phylogenetic analyses are listed in **S6 Table**. We used ggtree [50] and ete3 packages [51] for the visualization of trees.

### EBL classification according to current bornaviral genera

EBLs were classified into current bornaviral genera based on the query bornaviral sequence with the lowest E-value in the tBLASTn search. The results of the phylogenetic analysis-based method and the similarity score-based method were highly concordant (EBLN: 99.7%, EBLG: 100.0%, and EBLL: 100.0%). Thus, we applied the classification method for all EBL loci (**Fig 1C**). We could not create reliable phylogenetic trees using EBLP or EBLM due to the small number of sites available for phylogenetic analysis.

### Assessment of genetic diversity of ancient bornaviral sequences

We compared the genetic diversity of ancient and extant bornaviruses to infer how diverse bornaviruses have endogenized during primate evolution. First, we used the most recent common ancestor of the EBLN orthologs as the ancestral bornaviral N gene to avoid overestimating the sequence diversity of ancient bornaviruses. Next, to provide a comparison standard for interpreting the ancient bornaviral genetic diversity, we calculated the genetic distance for classifying extant bornaviral species in our phylogenetic tree (0.06 substitutions per site) (**S3 Table**). The genetic distances between nodes in the phylogenetic tree were calculated using the ete3 toolkit. It should be noted that this is an alternative method, and ICTV classification for extant bornaviral species is based on the sequence similarity among intact viral genomes, differences in their host ranges, and phylogenetic analysis using viral proteins.

## Data Availability

The codes are available at https://github.com/Junna-Kawasaki/EBL_2020. The versions of bioinformatics tools are listed in **S7 Table**.

## Acknowledgments

We thank Dr. Keiko Takemoto (Institute for Virus Research, Kyoto University, Japan) for technical support. We are grateful to Jumpei Ito (Institute of Medical Science, the University of Tokyo, Japan), Bea Clarise Garcia, Lin Hsien Hen, Koichi Kitao, and Michiko Iwata (Institute for Frontier Life and Medical Sciences, Kyoto University) for helpful discussions.

This study was supported by JSPS KAKENHI JP19J2224 (JK); and JP18K19443 (MH); MEXT KAKENHI JP17H05821 (MH) and JP19H04833 (MH); Hakubi project at Kyoto University (MH). Computations were partially performed on the supercomputing systems SHIROKANE (Human Genome Center, the Institute of Medical Science, the University of Tokyo) and the NIG supercomputer (ROIS National Institute of Genetics).

## Author contributions

MH conceived the study; JK mainly performed bioinformatics analyses; MH, SK, and YM supported bioinformatics analyses; JK prepared the figures and wrote the initial draft of the manuscript; all authors contributed designed the study, interpreted data, revised the paper, and approved the final manuscript.

## Supporting information

**S1 Fig. Dating analysis for bornaviral integration events. (A)** Procedure to determine presence and absence patterns of orthologous EBLs. First, we performed pairwise alignment among EBL integration sites using BLASTN and made an all-against-all matrix of their alignment coverages. Second, we constructed a network using the matrix and grouped EBL loci by extracting community structures. Finally, the ages of bornavirus integration events were assigned from the divergence times of host species with orthologous EBLs. **(B)** The all-against-all matrix of alignment coverages among EBL integration sites. In the heatmap, the blue color palette shows the alignment coverage between EBL integration sites (%) and yellow indicates that sequence similarity was not detected (ND). The column colors indicate EBL groups; in particular, the white shows manually modified groups (EBLG2, EBLL2, EBLL35, and EBLL36) (**details in Materials and Methods**). The row colors show host lineages of each EBL locus.

**S2 Fig. Alignment quality between EBL integration sites. (A-C)** Schematic images of alignments between EBL integration sites. The sequence alignments of EBLG2 (A), EBLL2 (B), and EBLL35 (C) were visualized using AliTV. Blue lines indicate host chromosomal DNA, and the location of EBLs are shown as white colored portions of the lines. The black vertical lines are shown for every 1000 bp. The color palette from red to green indicates identity scores obtained from lastz. The representative host species are shown as silhouettes to the left of the alignments. **(D)** Dot plot between laurasiatherian and primate EBLG2 integration sites. The line colors except for gray correspond to (A), and gray lines indicate short fragments that were aligned by lastz. The white portions within the blue thick lines indicate the positions of EBLG2 in the genomes.

**S3 Fig. Phylogenetic tree of EBLNs and modern bornaviral N proteins.** These trees were constructed by the maximum likelihood method using amino acid sequences of EBLN and modern bornaviral N genes of the genus Carbovirus (A), Orthobornavirus (B), or Cultervirus (C). Colored arrows indicate extant bornaviruses. The color of external nodes indicates the extant bornaviral genus or the host species in which the EBLN was identified, as shown in the lower right corner. The square or triangle labels on the internal nodes correspond to the collapsed nodes in **Fig 3A-C**. The colored boxes highlight the bornaviral lineages endogenized during primate evolution. The numbers on the branches are bootstrap values (%) based on 1000 replications. The scale bars show genetic distances (substitutions per site). The genetic distance to distinguish extant bornaviral species is shown as the comparative standard for estimating the genetic diversity of ancient bornaviruses.

**S4 Fig. Validation of the network-based method for detection of orthologs using human transposable elements. (A)** Strategy for evaluating the detection rate of orthologs using our network-based method. To validate our network-based method, we compared it with the method of detecting orthologs using genomic alignments. First, we estimated the integration age of all human transposable elements (TEs) by LiftOver using the genomic alignment among 18 mammalian species shown in (B). Second, we randomly sampled 100 loci for each of the nine timescales, shown as a to i in (B), from the dating results of the genomic alignment-based method. Using these test datasets, we performed dating analysis by our network-based method. Third, we compared the results between the two methods by checking the predicted ages and detected orthologs. Example 1: integration ages coincided between two methods, and our method could detect all orthologs defined by the genomic alignment-based method. Example 2: integration ages coincided between two methods, but our method detected an incomplete set of orthologs. Example 3: the integration ages were mismatched between the two methods. Example 4: estimation ages were matched between two methods, but there was a contamination of sequence unrelated to true orthologous relationships. Furthermore, to select the best criteria for our network-based dating analysis, we tried nine different criteria shown in (C) (**details in Materials and Methods**). **(B)** Phylogenetic tree of mammalian species used to detect orthologs of human TEs. These 18 mammalian species were used to detect orthologs of human TEs based on the genomic alignment obtained from the UCSC genome browser. To validate the network-based dating method for each timescale, we randomly sampled 100 TE loci from nine different timescales (a to i). **(C)** Concordant rates between two methods for estimating integration ages. Each panel shows the result using different criteria for network construction (**details in Materials and Methods**). The x-axis indicates the timescales shown in (B). The y-axis indicates the concordant rate (%) between estimations using the genomic alignment-based method and our network analysis-based method. The blue labels indicate the concordant rates (%) between the two methods at the criteria used for the dating analysis for EBLs.

**S1 Table. The genomic position of EBLs**

**S2 Table. Reference list for mammalian biogeography related to ancient bornaviral infections**

**S3 Table. Genetic distances between ancient and extant bornaviral N genes**

**S4 Table. Accession numbers of bornaviral sequences used for the tBLASTn search**

**S5 Table. Chain files and genome assemblies used for validation for our dating method**

**S6 Table. Extant viral sequences used for phylogenetic analyses**

**S7 Table. Bioinformatics tools used in this study**

