## Supplementary figures and images for "One hundred million years history of bornavirus infections hidden in vertebrate genomes"

### Supplementary Figures

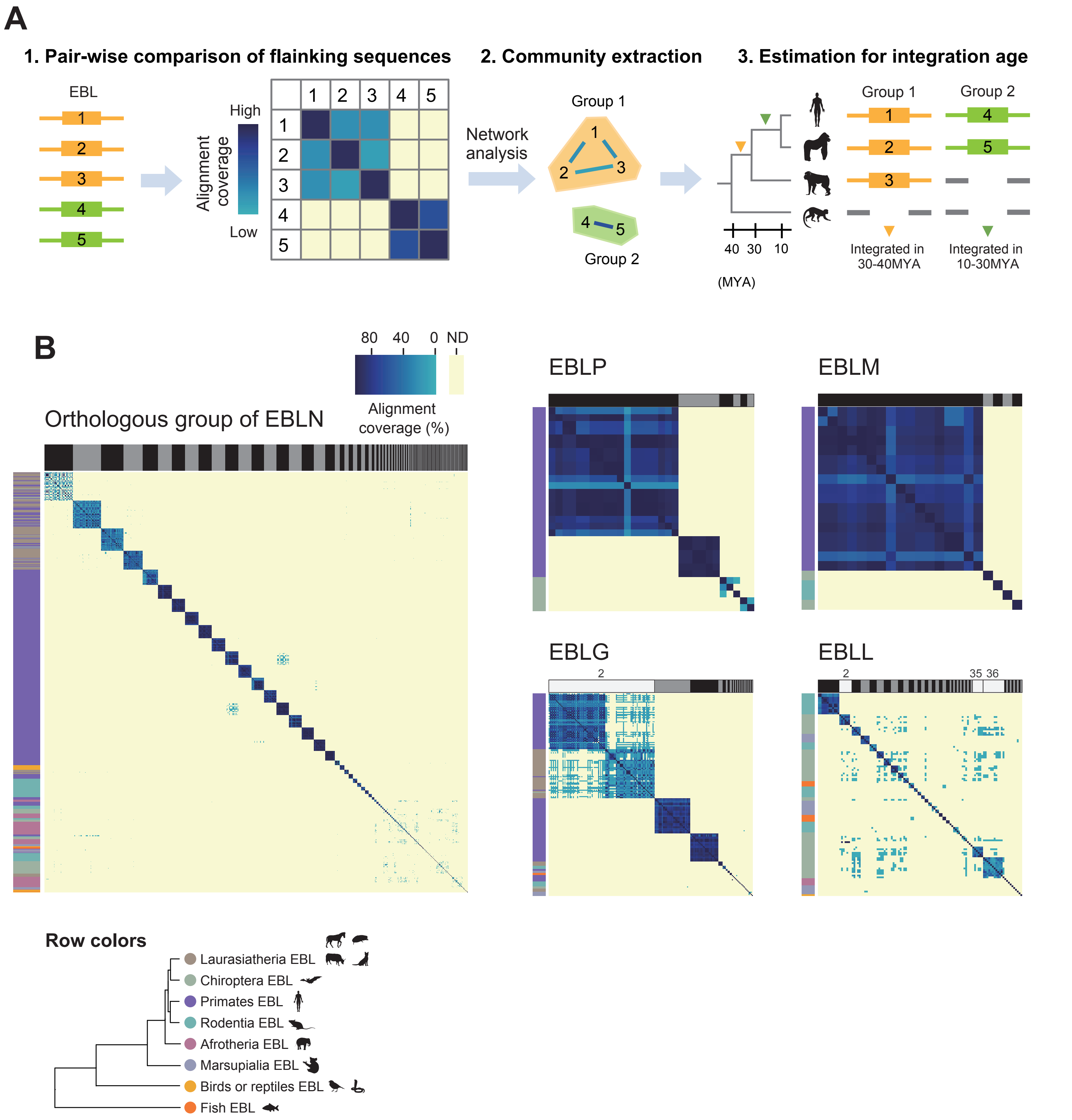


Supplementary Figure 1

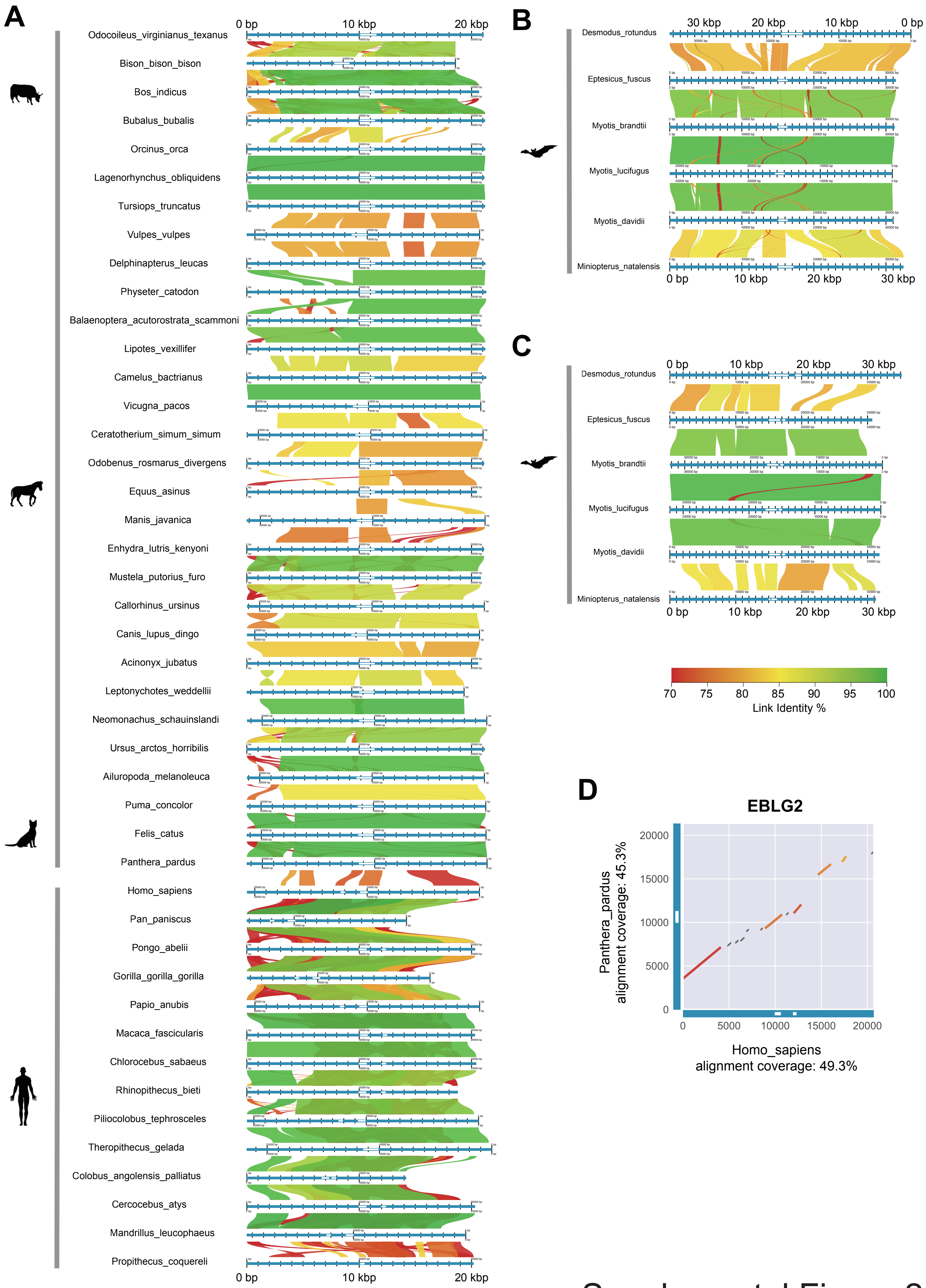


Supplemental Figure 2

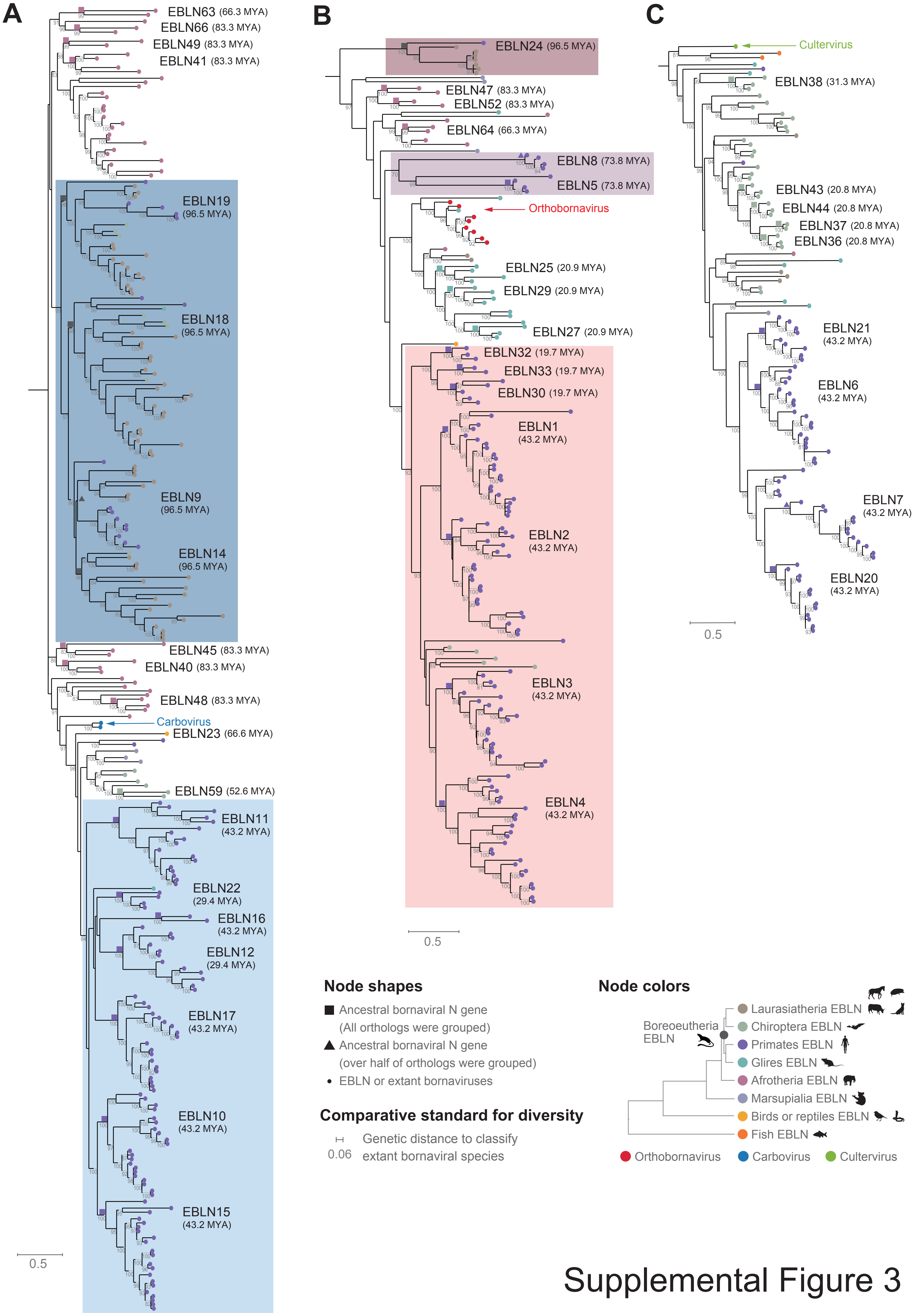


Supplemental Figure 3

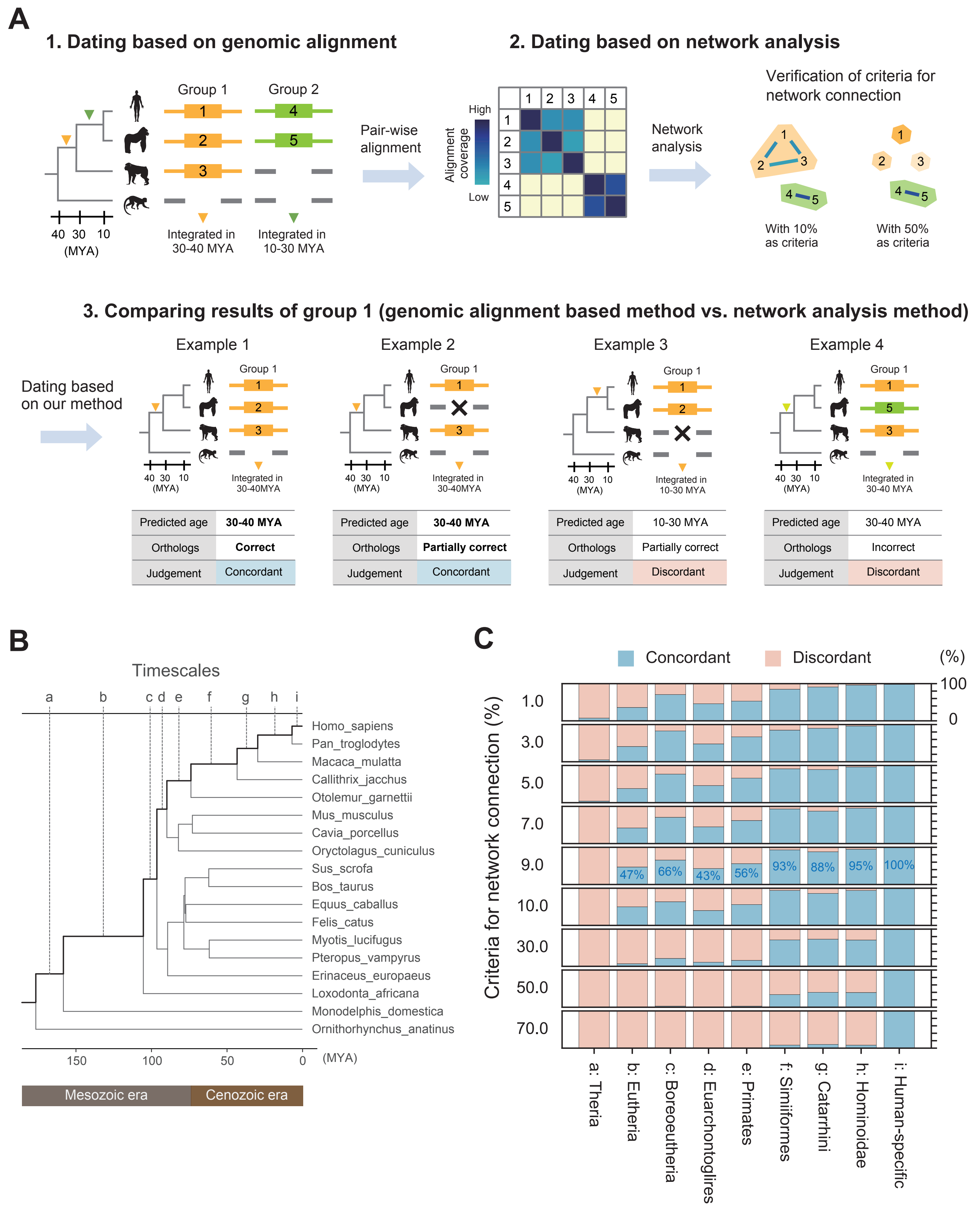


Supplemental Figure 4
